# Large-scale whole-genome sequencing of three diverse Asian populations in Singapore

**DOI:** 10.1101/390070

**Authors:** Degang Wu, Jinzhuang Dou, Xiaoran Chai, Claire Bellis, Andreas Wilm, Chih Chuan Shih, Wendy Wei Jia Soon, Nicolas Bertin, Chiea Chuen Khor, Michael DeGiorgio, Sonia Maria Davila Dominguez, Patrick Tan, Asim Shabbir, Angela Moh, Eng-King Tan, Jia Nee Foo, Tan Tock Seng Hospital Healthy Control Workgroup, Roger S. Foo, Carolyn S.P. Lam, A. Mark Richards, Ching-Yu Cheng, Tin Aung, Tien Yin Wong, Jianjun Liu, Chaolong Wang, on behalf of the SG10K Consortium

## Abstract

Asian populations are currently underrepresented in human genetics research. Here we present whole-genome sequencing data of 4,810 Singaporeans from three diverse ethnic groups: 2,780 Chinese, 903 Malays, and 1,127 Indians. Despite a medium depth of 13.7×, we achieved essentially perfect (>99.8%) sensitivity and accuracy for detecting common variants and good sensitivity (>89%) for detecting extremely rare variants with <0.1% allele frequency. We found 89.2 million single-nucleotide polymorphisms (SNPs) and 9.1 million small insertions and deletions (INDELs), more than half of which have not been cataloged in dbSNP. In particular, we found 126 common deleterious mutations (MAF>0.01) that were absent in the existing public databases, highlighting the importance of local population reference for genetic diagnosis. We describe fine-scale genetic structure of Singapore populations and their relationship to worldwide populations from the 1000 Genomes Project. In addition to revealing noticeable amounts of admixture among three Singapore populations and a Malay-related novel ancestry component that has not been captured by the 1000 Genomes Project, our analysis also identified some fine-scale features of genetic structure consistent with two waves of prehistoric migration from south China to Southeast Asia. Finally, we demonstrate that our data can substantially improve genotype imputation not only for Singapore populations, but also for populations across Asia and Oceania. These results highlight the genetic diversity in Singapore and the potential impacts of our data as a resource to empower human genetics discovery in a broad geographic region.

## Introduction

We have learned profound insights into the demographic history of humans and the genetic basis of phenotype diversity and disease etiology by mining the variation of human genomes.^1-4^ While earlier efforts were mostly based on array genotyping of common variants, large-scale whole-genome sequencing (WGS) has become a powerful approach for disease gene mapping by surveying the genome in an unbiased and completed fashion. A remarkable milestone is the completion of the 1000 Genomes Project (Phase 3, 1KG3), which sequenced >2,500 genomes from 26 populations globally at ~7.4× and cataloged over 88 million variants segregated in the sample.^5^ The resource provided by 1KG3 has empowered numerous genetic studies through imputation of large numbers of samples genotyped in genome-wide association studies (GWAS), allowing for detection of genetic association at low frequency variants (minor allele frequency, MAF<0.05) and thus enabling a deeper understanding of the genetic architecture of complex diseases.^6,7^ The direct assessment of causal variants enabled by sequencing technologies has led to the convergence of population and clinical genetics.^8,9^ Population genetic information has become the cornerstone of precision medicine, with wide applications in the diagnosis of Mendelian diseases,^10,11^ optimization of therapeutic treatments,^12^ drug development,^13^ and disease risk prediction and stratification.^14^

Comparing across populations globally, the majority of single-nucleotide variants are rare (MAF<0.005) and population-specific, highlighting the importance of population diversity in human genomics research.^5,11,15^ To implement precision medicine for local populations, many countries have initiated population-based WGS studies.^16-19^ Nevertheless, current public WGS resources are predominantly based on Europeans. Though Asia is the largest continent with over four billion inhabitants from diverse populations,^4^ Asian genomes are underrepresented in public databases. For example, the Haplotype Reference Consortium (HRC) has brought together WGS data of 32,488 samples from 20 different studies to construct a large reference panel for accurate imputation.^20^ This HRC panel, however, is only appropriate for imputing European subjects. Moreover, in the genome Aggregation Database (gnomAD), only ~5% of its 15,496 genomes are from East Asians and none are from South Asians. Furthermore, in the Trans-Omics for Precision Medicine (TOPMed) Program, one of the largest ongoing WGS efforts supported by the United States National Heart, Lung, and Blood Institute (NHLBI), only ~7% of its >72,000 samples (Phases 1 and 2) are of Asian ethnicity (ASHG 2016, presentation by Gonçalo Abecasis).

Despite its small geographic size, Singapore has a diverse population-genetic composition due to its migratory history.^21-23^ Singaporeans are broadly classified into four ethnic groups, namely Chinese, Malay, Indian and other (CMIO). Each ethnic group further harbors substantial fine-scale genetic diversity. According to the Singapore Department of Statistics (latest update on 2017), Chinese account for 74.3% of the four million Singapore residents. Most Chinese are descendants of several dialect groups from south China and a minority are from north China. Malays, representing 13.4% of the Singapore population, include descendants of diverse Austronesian-speaking groups in Southeast Asia, primarily from Singapore, Malaysia, and Indonesia. About 9.1% of the population are Indians descending from the Indian migrants during the period of British colonization. The majority of Indians are Telugas and Tamils from southeastern India and a minority are Sikhs and Pathans from north India.^21^ The remaining 3.2% of the population are mainly Eurasians from Europe and Middle East. Taken together, the three major ethnicity groups in Singapore provide a unique snapshot of the genetic diversity across East Asia, Southeast Asia, and South Asia. Therefore, WGS analysis of Singaporeans have the potential to benefit a large number of Asian populations.

We have therefore initiated the SG10K project to whole-genome sequence 10,000 healthy individuals, as well as patients from several ongoing disease studies in Singapore. Our main objectives are to (1) comprehensively characterize genetic variation in Singapore populations; (2) create a WGS reference panel for accurate genotype imputation in Asian populations; and (3) generate a common control dataset to empower disease association studies. We chose a medium coverage design of ~15× to maximize the statistical power for rare variant association studies given a fixed sequencing budget.^24,25^ In this article, we describe the Phase 1 data of SG10K and initial findings based on 4,810 whole genomes, including 2,706 healthy samples without major diseases and 2,104 patient samples with heart failure, Parkinson’s disease, and obesity. We characterize genetic mutations segregating in the population and in personal genomes and discuss the implications of our findings for precision medicine. Furthermore, we investigate the genetic diversity and fine-scale population structure in Singapore, by comparing to worldwide populations from the 1KG3 dataset.^5^ Lastly, we illustrate the potential impacts of the SG10K data in improving imputation accuracy for both Singapore populations and worldwide populations from the Human Genome Diversity Project (HGDP).^26^

## Results

### Dataset overview

Phase 1 of the SG10K Project includes 4,810 samples successfully sequenced at a mean depth of 13.7×. These samples were contributed by eight cohorts in Singapore, including five healthy cohorts free of major diseases and three patient cohorts of heart failure, Parkinson disease, and obesity (**Supplementary Table 1**). Before genotype calling, we inferred ethnicity, sex, and contamination rate for each individual directly from sequence reads (**Materials and Methods, Supplementary Figures 1-4, Supplementary Tables 2-3**).^27-29^ We obtained concordance rates of 96.7% between inferred and self-reported ethnicity, and 99.2% between inferred and self-reported sex. This lower ethnicity concordance rate was mostly due to admixture between different ethnic groups. Based on the inferred ethnicity or sex, we classified 4,810 samples into 2,780 Chinese, 903 Malays, and 1,127 Indians, as well as 2,462 females and 2,348 males. We performed linkage-disequilibrium (LD) based joint calling for all SG10K samples, followed by a series of quality controls to filter low-quality variants (**Supplementary Figure 1**).^30^ Notably, we inferred 494 pairs of close relatives (third degree or closer), including 93 duplicates (or monozygotic twins), 17 trios, and 32 parents-offspring duos (**Supplementary Figure 5, Supplementary Table 4**).^31,32^ Sixteen out of the 17 trios were from the Platinum Asian Genomes Project, which recruited 46 individuals from 14 trios and one quartet family (two siblings and their parents). Seventy-two out of 93 duplicates were subsequently confirmed to be heart failure patients recruited by different studies at different time points. The majority of the remainder exhibited previously unknown cryptic relatedness, including 123 cross-cohort pairs. By removing close relatives up to the third degree, we obtained a maximum unrelated subset consisting of 4,446 individuals, 2,535 of which were healthy individuals. We utilized the duplicates, trios, and duos to further filter out low-quality variants with more than two duplicate or Mendelian discordance instances. Lastly, we applied a population-based phasing algorithm to phase genotypes to haplotypes for all samples including close relatives.^33^ The final dataset consisted of haplotypes from 4,810 samples, including 89,160,286 SNPs and 9,113,420 INDELs from 22 autosomes and the X chromosome. We detected slightly more SNPs and many more INDELs compared to those reported by the 1KG3 (84.7 million SNPs and 3.6 million INDELs),^5^ which is likely attributed to our larger sample size and higher sequencing depth than 1KG3.

**Table 1.** Evaluation of final SG10K call set based on array genotyping data of 1,263 samples on chromosome 2.

|  | Minor allele frequency (MAF) bins |  |  |  |
| --- | --- | --- | --- | --- |
|  | 0.01-0.05 | 0.05-0.2 | 0.2-0.5 | All |
| Number of SNPs on array | 1,070 | 13,208 | 25,770 | 40,048 |
| Sensitivity | 0.9981 | 0.9990 | 0.9973 | 0.9979 |
| Non-reference sensitivity | 0.9991 | 0.9996 | 0.9997 | 0.9997 |
| Precision | 0.9976 | 0.9984 | 0.9981 | 0.9982 |
| Overall concordance | 0.9995 | 0.9996 | 0.9994 | 0.9995 |
| Heterozygote concordance | 0.9973 | 0.9990 | 0.9992 | 0.9992 |
| Non-reference concordance | 0.9979 | 0.9992 | 0.9993 | 0.9993 |

Definitions: sensitivity, fraction of polymorphic sites in the array data that were also in the sequencing call set with both alleles matched; non-reference sensitivity, fraction of non-reference genotypes in array data that are also called as non-reference genotypes in sequencing data; precision, fraction of concordant genotypes across non-reference sites in the sequencing call set; overall/heterozygote/non-reference concordance, fraction of concordant genotypes across all/heterozygous/non-reference sites in array data.

**Table 2.**
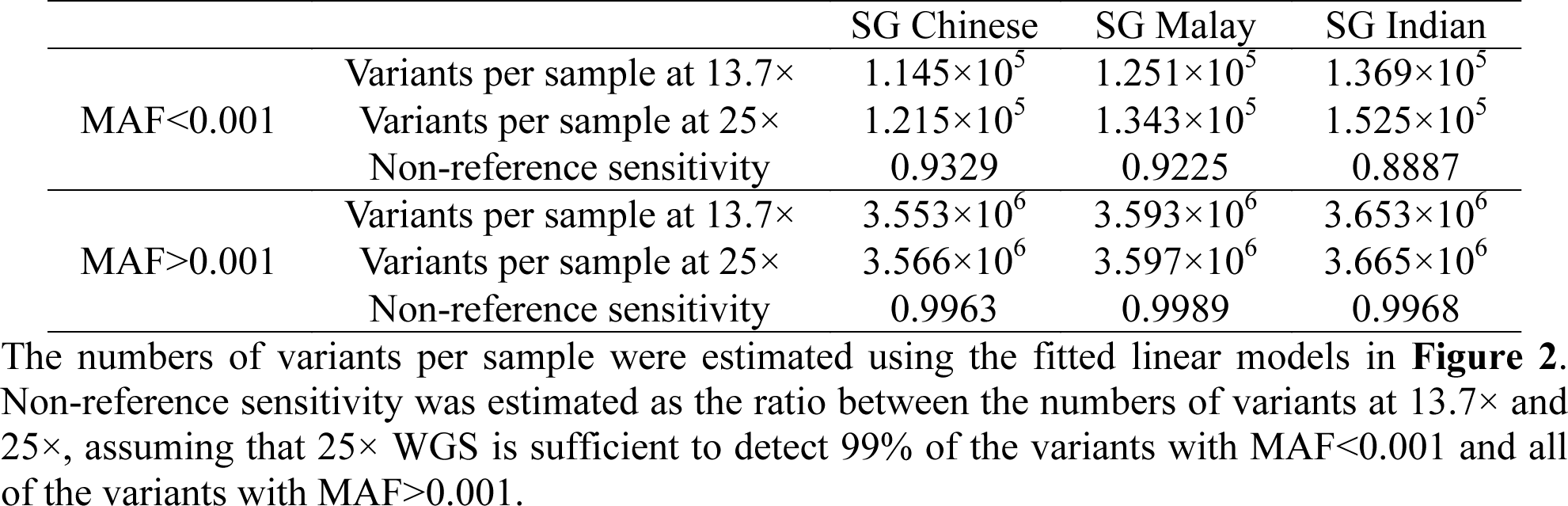
Estimated sensitivity to detect variants with minor allele frequency below and above 0.001 in the SG10K call set for different ethnicity groups.

|  |  | SG Chinese | SG Malay | SG Indian |
| --- | --- | --- | --- | --- |
| MAF<0.001 | Variants per sample at 13.7× | $1.145 \times 10^5$ | $1.251 \times 10^5$ | $1.369 \times 10^5$ |
| | Variants per sample at 25× | $1.215 \times 10^5$ | $1.343 \times 10^5$ | $1.525 \times 10^5$ |
|  | Non-reference sensitivity | 0.9329 | 0.9225 | 0.8887 |
| MAF>0.001 | Variants per sample at 13.7× | $3.553 \times 10^6$ | $3.593 \times 10^6$ | $3.653 \times 10^6$ |
| | Variants per sample at 25× | $3.566 \times 10^6$ | $3.597 \times 10^6$ | $3.665 \times 10^6$ |
|  | Non-reference sensitivity | 0.9963 | 0.9989 | 0.9968 |

**Figure 1.**
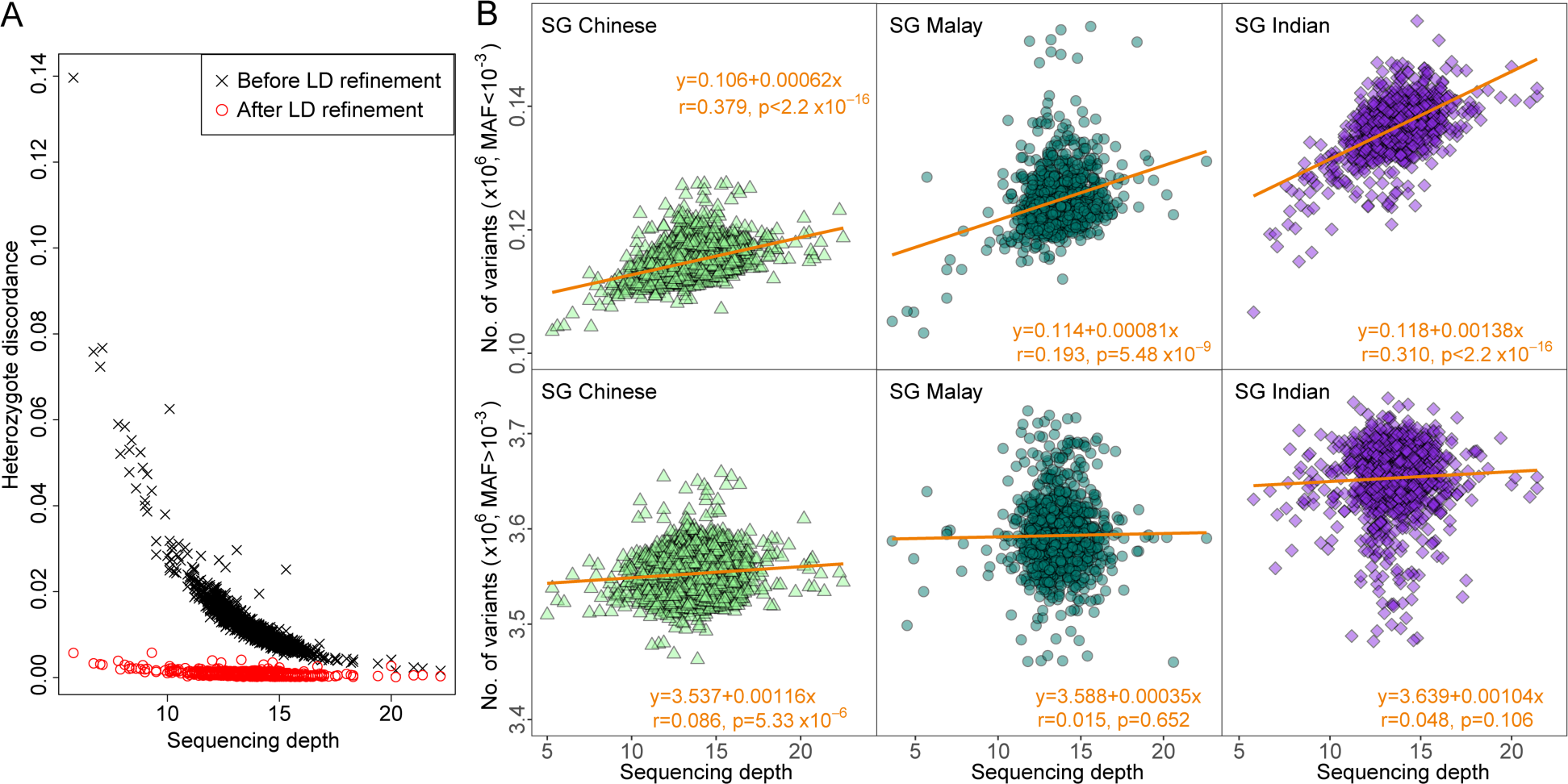
Quality evaluation of the SG10K call set. (A) Heterozygote discordance rate versus sequencing depth for 1,263 samples that have array genotyping data. Discordance rate was computed based on 39,964 SNPs on chromosome 2 that were in both sequencing call set and array data set. (B) Number of variants detected in each sample as a function of sequencing depth. In each of subpanels of (B), outliers more than 5 standard deviations from the mean sequencing depth (x-axis) or the mean number of variants (y-axis) were removed and not shown. A linear model (orange line and text) was fitted based on the remaining samples. The *p* value was derived from the Student’s *t* test against the null hypothesis of Pearson correlation *r*=0. Results without removing outliers are shown in **Supplementary Figure 6**.

### Quality evaluation

We estimated a transition to transversion ratio (Ts/Tv) of 2.07 across 84.7 million bi-allelic autosomal SNPs in our final dataset, which is consistent with values reported by previous studies.^5,17^ We further evaluated the quality of our call set based on 1,263 individuals previously genotyped by the Illumina Quad610 array.^34^ Treating the array data as gold standard, we achieved a 0.9997 non-reference sensitivity and a 0.9992 heterozygote concordance rate for variants with MAF>0.01, despite the medium coverage design (**Table 1**). For low-frequency variants with MAF between 0.01 and 0.05, both the sensitivity and heterozygote concordance rate dropped slightly to 0.9991 and 0.9973, respectively. These statistics indicate high quality of our call set, especially due to the proper use of shared LD among a large number of samples. In particular, before LD-based refinement, the discordance rate of each sample highly depended on the sequencing depth and was as high as ~0.075 at ~7× (**Figure 1A**). However, after LD-based refinement, the discordance rate was reduced dramatically by 25-fold to ~0.003 for samples sequenced at ~7×, and by 10-fold from ~0.01 to ~0.001 for the majority of samples sequenced at a medium depth of ~13.7× (**Figure 1A**).

A primary issue of sequencing at a medium depth is the sensitivity to detect rare variants, which cannot be evaluated by comparing to GWAS array data. We thus designed an approach to estimate sensitivity without external data for comparison. For each sample, we counted the number of non-reference variants at extremely low frequency (MAF<0.001). Within each population, we observed that samples sequenced at lower depths carried fewer rare variants (Pearson’s r>0.2, p<10^-8^, Student’s *t* test), indicating lower sensitivity at detecting rare variants (**Figure 1B**). In contrast, the mutation burden did not have an obvious trend with sequencing depth for variants with MAF>0.001 (Pearson’s r<0.07, **Figure 1B**). Both simulation studies^24^ and our empirical data (**Supplementary Figure 6**) suggested the power to detect extremely rare variants plateaus at ~25×. By assuming a linear model and a 0.99 non-reference sensitivity at 25× sequencing depth, we estimated 0.9329, 0.9225, and 0.8887 non-reference sensitivities at detecting variants with MAF<0.001 in our data for Chinese, Malays, and Indians, respectively (**Table 2**). For variants with MAF>0.001, assuming 25× sequencing depth has perfect power, we estimated non-reference sensitivities to be 0.9963, 0.9989, and 0.9968 for Chinese, Malays and Indians in our call set, respectively, which is slightly lower than the non-reference sensitivity estimated by direct comparison with GWAS data for variants at a higher frequency of MAF>0.01 (**Table 1**). Compared to Chinese and Malays, the lower sensitivity at detecting rare variants in Indians might be attributed to their higher genetic diversity, consistent with the greater number of non-reference variants per Indian genome (**Figure 1B**).

### Novel variants and implications for genetic diagnosis

In our final dataset, 45.6 million SNPs (51%) and 6.3 million INDELs (70%) were novel variants not included in dbSNP (version 150). This higher proportion of novel INDELs might be partially attributed to multiple representations and ambiguous positions of the same INDELs in previous studies, and here we used the parsimonious left-aligned representation to normalize INDELs.^35^ We also found that chromosome X has higher proportion of novel variants than the autosomes consistently across different variant types (**Supplementary Table 5**), suggesting that chromosome X has been less studied than autosomes. Unsurprisingly, the majority of our novel variants are extremely rare (**Supplementary Figure 7A**), consistent with previous studies.^5^ Specifically, singletons and doubletons accounted for 69.6% and 14.4% of the novel SNPs, respectively, which are much higher than those of the known SNPs. Moreover, only ~0.5% of the novel variants reached MAF>0.01 (equivalent to minor allele counts MAC>90).

Among the common (MAF>0.01) novel variants, we identified 126 “deleterious” mutations annotated by Polyphen or SIFT.^36-38^ These variants were mapped to 113 genes, including 99 genes reportedly harboring at least one pathogenic or likely pathogenic variants according to ClinVar (**Supplementary Figure 7B**).^39^ However, the high frequencies of these variants in local populations suggest that they are likely benign or have very low penetrance.^40^ Comparing across Chinese, Malays and Indians, none of the 126 “deleterious” mutations were population-specific, whereas 35 reached MAF>0.05 in all populations (**Supplementary Figure 6C**), indicating a high risk of genetic misdiagnoses if Singaporean patient genomes were compared to existing public databases where these variants are currently absent.^11^

### Variants in a personal genome

To compare typical genomes between different ethnic groups, we focused on 2,535 individuals free of major diseases (**Supplementary Table 6**). We found an average healthy individual carries autosomal variants at ~3,330,000 SNPs, ~105,000 insertions, and 160,000 deletions (**Table 3**). Across putatively functional categories, we observed that a genome harbors ~11,600 missense variants, ~54,000 variants overlapping with untranslated regions (UTRs), ~5,000 variants at transcription factor binding sites (TFBS), ~2,000 deleterious variants predicted by SIFT or Polyphen, and ~31 pathogenic variants as annotated by ClinVar. Moreover, Indian genomes possessed the highest number of variants on average, followed by Malays and Chinese. In addition, the heterozygote to non-reference homozygote ratio (Het/Hom) was also higher in Indian (1.73) than in Chinese (1.43) and Malays (1.51), reflecting a higher level of genetic diversity among Indians. Though Malays have a lower level of genetic diversity than Indians,^21^ sequencing a Malay genome leads to a discovery of ~30.2 thousand novel SNPs, which is close to ~30.3 thousand for an Indian and higher than ~27.6 thousand for a Chinese, which reflects the more severe underrepresentation of Southeast Asians in current genetic studies. We note that the Het/Hom ratio was high for novel variants (~13 for SNPs and ~25 for INDELs) because the majority of the novel variants are rare and often present as heterozygotes. Furthermore, except for novel variants, the highest Het/Hom ratio was among variants annotated as deleterious or pathogenic, which is consistent with negative selection against these variants to reach high frequencies.

**Table 3.** The median number of autosomal variant per genome in Singapore populations.

| Variant annotation | Singapore Chinese<br>N=1,267, depth=13.7× |  | Singapore Malay<br>N=454, depth=13.8× |  | Singapore Indian<br>N=814, depth=13.6× |  |
| --- | --- | --- | --- | --- | --- | --- |
|  | Variants | Het/Hom | Variants | Het/Hom | Variants | Het/Hom |
| SNP | 3,308,882 | 1.43 | 3,346,583 | 1.51 | 3,406,391 | 1.73 |
| Insertion | 103,206 | 1.50 | 106,398 | 1.61 | 116,305 | 1.91 |
| Deletion | 158,864 | 1.85 | 162,288 | 1.98 | 170,818 | 2.33 |
| SNP not in dbSNP150 | 27,635 | 12.52 | 30,155 | 13.55 | 30,263 | 14.02 |
| Insertion not in dbSNP150 | 6,527 | 23.16 | 7,089 | 24.49 | 8,928 | 25.53 |
| Deletion not in dbSNP150 | 6,059 | 25.55 | 6,478 | 26.08 | 8,058 | 25.79 |
| Synonymous | 11,675 | 1.45 | 11,844 | 1.55 | 12,077 | 1.78 |
| Missense | 11,509 | 1.51 | 11,666 | 1.60 | 11,911 | 1.81 |
| Exon | 127,268 | 1.46 | 128,961 | 1.56 | 131,579 | 1.78 |
| Intron | 1,859,670 | 1.46 | 1,884,753 | 1.55 | 1,925,723 | 1.77 |
| UTR | 53,751 | 1.46 | 54,418 | 1.55 | 55,564 | 1.79 |
| TFBS | 4,911 | 1.61 | 4,995 | 1.72 | 5,141 | 1.99 |
| SIFT: Deleterious | 1,672 | 2.81 | 1,697 | 3.00 | 1,715 | 3.48 |
| Polyphen: Probably damaging | 877 | 3.34 | 896 | 3.56 | 902 | 4.15 |
| Polyphen: Possibly damaging | 1,150 | 2.92 | 1,165 | 3.04 | 1,182 | 3.52 |
| ClinVar: Pathogenic | 30 | 2.33 | 31 | 2.67 | 33 | 3.38 |
| ClinVar: Association | 46 | 1.80 | 45 | 2.14 | 43 | 2.28 |
| ClinVar: Risk factor | 82 | 1.45 | 82 | 1.61 | 82 | 1.89 |

Only healthy individuals without any known major diseases were included in this analysis. Abbreviations: UTR, untranslated region; TFBS, transcription factor binding site; Het/Hom, ratio of number of heterozygotes to non-reference homozygotes. Annotations were from the Variant Effect Predictor (VEP-v91). For ClinVar annotations, “pathogenic” variants are those interpreted for Mendelian disorders by the American College of Medical Genetics and Genomics and the Association for Molecular Pathology (ACMG/AMP); “association” variants are those identified by GWASs and further interpreted for their clinical significance; “risk factor” variants are those interpreted not to cause a disorder but to increase the disease risk.

The ClinVar pathogenic variants are interpreted for Mendelian disorders and might cause adverse clinical outcome if present as homozygous, especially for recessive disorders. We found that even without any known major diseases, each individual carried 3.9±2.0 (mean±s.d.), 4.3±2.1, and 4.9±2.2 pathogenic homozygotes in Chinese, Malays, and Indians, respectively. Moreover, individuals with higher inbreeding coefficients tended to have a greater number of pathogenic homozygotes (**Supplementary Figure 8**), although such a trend might be underestimated because our analysis was restricted to healthy individuals and we did not recruit patients with Mendelian disorders. In this analysis, we also noticed that Indians had a long tail distribution of inbreeding coefficients, which is consistent with a high level of consanguinity in Dravidian south India and Pakistan.^41^ We estimated the prevalence of consanguineous mating between second cousins or closer relatives (inbreeding coefficient >0.0156) to be 29.1% (237/814) in Indians, followed by 10.8% (50/464) in Malays, and 2.6% (33/1267) in Chinese. These results are important, as consanguineous mating can lead to an excess of infant/childhood mortality and extended morbidity.^41^

### Population structure and genetic diversity

We analyzed Singapore populations together with 26 worldwide populations from the 1KG3 dataset using principal components analysis (PCA). To avoid undesirable effects on PCA due to uneven sample sizes,^42^ we randomly downsampled each Singapore population to 100 unrelated individuals in the joint analysis with the 1KG3 populations, and subsequently projected the remaining SG10K samples into the PCA space.^28,29^ We found that SG Indians and SG Chinese overlapped largely with South and East Asians, respectively, whereas SG Malays formed a cluster distinct from any 1KG3 population (**Figure 2A**, **Supplementary Figure 9**). When restricting PCA to East/Southeast Asians, the majority of SG Chinese overlapped with CHS from south China and the rest with CHB from north China (**Supplementary Figure 10**). Similarly, we observed that the majority of SG Indians overlapped with STU, ITU, and BEB from the south of the Indian subcontinent, with a small proportion overlapping with GIH from west India and PJH from Pakistan (**Supplementary Figure 11**). Given the clear north-south pattern on both PCA of East Asians and South Asians, we estimated that 96% and 4% of the SG Chinese samples are from south and north China respectively, and that 81% and 19% of the SG Indian samples are from south and north India subcontinent respectively (**Materials and Methods**). In addition, neither SG Chinese nor Malays overlapped with CDX and KHV from the mainland Southeast Asia. These PCA results are consistent with the genetic distances between pairwise populations (**Supplementary Figure 12**), in which the SG Malay is relatively distant from the other populations with the closest being KHV (F_ST_=0.007) and CDX (F_ST_=0.009). Finally, we applied PCA on only the SG10K samples and found that three distinct clusters consisting of Chinese, Malays, and Indians emerged, with a noticeable number of likely admixed individuals forming clines between clusters (**Supplementary Figure 13**).

**Figure 2.**
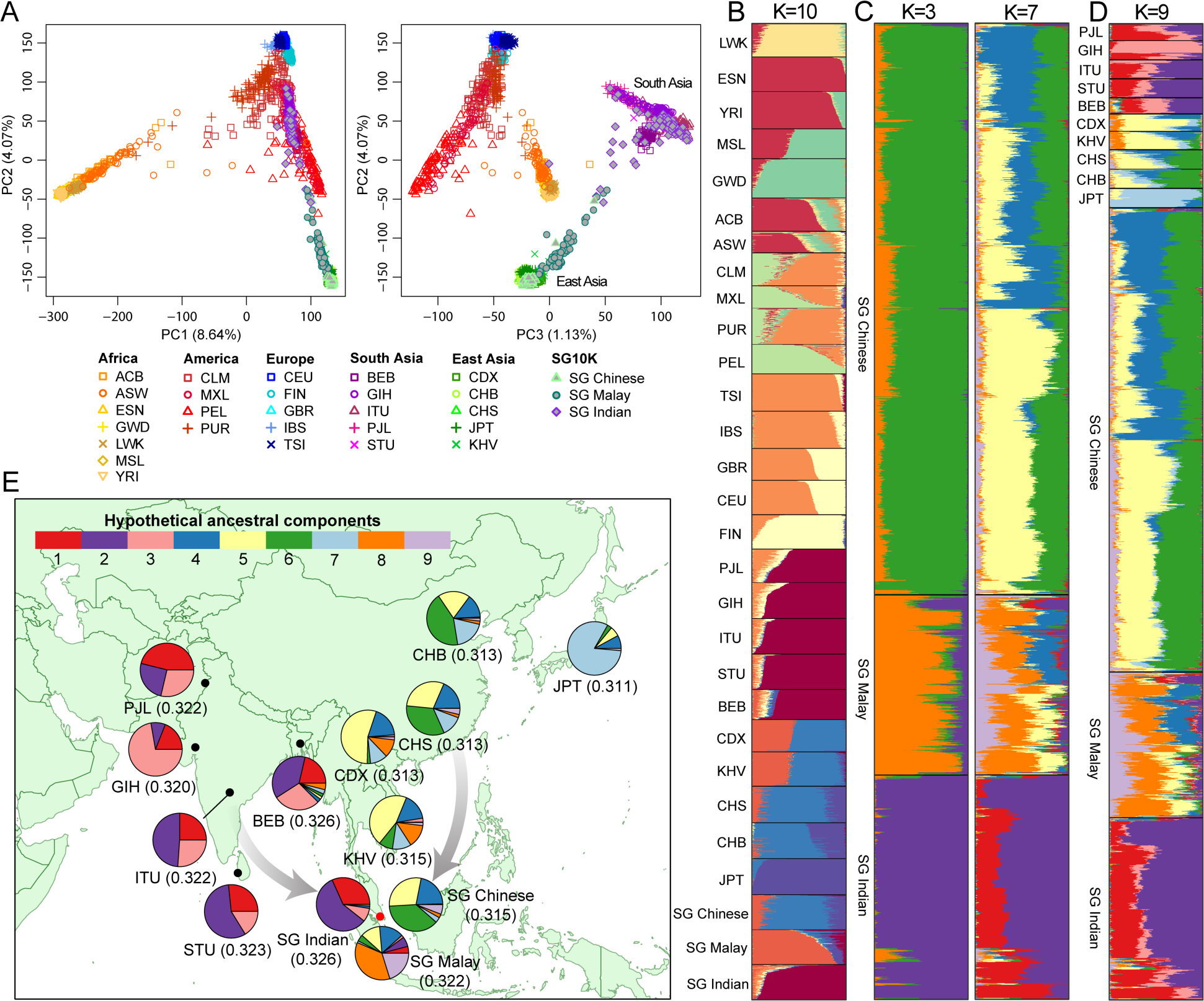
Population structure of SG10K and 1KG3 samples. (A) PCA of the SG10K and 1KG3 data. Proportion of variance explained by each PC is indicated in the axis label. (B) ADMIXTURE analysis of the SG10K and 1KG3 data (K=10). Each colored bar represents one individual and the length of each colored segment represents admixture proportion of an ancestral component. 100 unrelated individuals from each Singapore population were included in the analyses of (A) and (B). (C) ADMIXTURE analysis of 4,446 unrelated individuals from SG10K (K=3 and K=7). (D) ADMIXTURE analysis of 4,446 unrelated SG10K individuals together with South Asians and East Asians from 1KG3 (K=9). (E) Geographic distribution of the nine ancestry components in (D). Each pie chart represents the ancestry proportions averaged across individuals from the same population. Heterozygosity for each population was shown in the parentheses. Populations in 1KG3: ACB, African Caribbean; ASW, African American; ESN, Esan; GWD, Gambian; LWK, Luhya; MSL, Mende; YRI, Yoruba; CLM, Colombian; MXL, Mexican; PEL, Peruvian; PUR, Puerto Rican; CEU, Northern and Western European; FIN, Finnish; GBR, British; IBS, Iberian; TSI, Toscani; BEB, Bengali; GIH, Gujarati; ITU, Telugu; PJL, Punjabi; STU, Sri Lankan Tamil; CDX, Chinese Dai; CHB, Han Chinese in Beijing; CHS, Southern Han Chinese; JPT, Japanese; KHV, Kinh.

We further investigated population structure using ADMIXTURE, a maximum likelihood method that models each genome as a mixture of K hypothetical ancestral components.^43^ The value of K was chosen based on five-fold cross validation to optimize prediction of genotypes. When applying ADMIXTURE to the full 1KG3 dataset and 300 SG individuals, we inferred the optimal number of ancestral components to be K=10 (**Supplementary Figure 14**). SG Malays contributed a new component (colored by dark orange in **Figure 2B**), which was also present at moderate levels in KHV and CDX. It is worth noting that this Malay component appeared at low levels in all Han Chinese populations but was significantly higher in SG Chinese (0.148±0.006, mean±s.d.) than in CHB (0.034±0.005; p<10^-16^, Wilcoxon rank-sum test) and CHS (0.112±0.005; p<10^-5^, Wilcoxon rank-sum test), suggesting recent gene flow from Malays to Chinese in Singapore. When we applied ADMIXTURE to the set of 4,446 unrelated SG10K samples, K=3 effectively distinguished the three major ethnic groups with admixed samples apparent in each group (**Figure 2C**). However, we inferred the optimal number of components to be K=7, indicating fine-scale population structure within major ethnic groups (**Supplementary Figure 15**). Interestingly, a component colored by blue showed up in about half of the Chinese samples and half of the Malay samples from K=6 (**Supplementary Figure 15**). Given that most SG Chinese trace their origin to the Fujian and Guangdong provinces in south China, we first speculated that this blue component might distinguish different Chinese origins at the province level. However, we excluded this possibility after examining 103 samples with detailed records of origin, because no difference was found in the levels of the blue component between samples from different provinces (**Supplementary Figure 16**). Furthermore, we also excluded the possibility of artifacts caused by batch effects or health status, as these 103 samples were from the same healthy cohort, and were processed and sequenced using the same protocol and platform.

We next combined the SG10K samples with those from other Asian populations in the 1KG3 to investigate the geographic distribution of inferred ancestral components. We set K=9 in this combined analysis, which mimics the result of K=7 in the analysis of SG10K alone, with two additional components introduced by Gujarati and Japanese (**Figure 2D**). We numerically labeled these nine ancestral components as shown in **Figure 2E**. We applied hierarchical clustering based on F_ST_ relating these nine components and observed three major clusters, representing South Asian ancestry (components 1, 2 and 3), East Asian ancestry (components 4, 5, 6 and 7), and Southeast Asian ancestry (components 8 and 9, **Supplementary Figure 17**). The presence of multiple ancestral components within the same ancestry group is suggestive of fine-scale population structure. Specifically, the SG Indians were predominated by components 1 and 2, which together reflected a clear south-north cline among SG Indians (**Figure 2D-E**). We found that component 3 introduced by the Gujarati Indians of 1KG3 contributed little to the SG Indians. Interestingly, we observed that the mysterious blue component described previously, which is now component 4, is prevalent across East and Southeast Asian populations—however, the origin of this component remained unexplained. We also noticed that component 4 was closely related to component 5 (F_ST_=0.007), which was mostly found in CDX in southwestern China and gradually declined toward both the north and the south. In contrast, we saw that components 6 and 7, which were mostly found in northern Chinese and Japanese respectively, had little contribution to SG Malays. Instead, the two major components in SG Malays were components 8 and 9, both reflecting indigenous Southeast Asian ancestry. In particular, component 8 was also present at moderate levels in mainland Southeast Asians (KHV and CDX) whereas component 9 was specific to SG Malays. In addition to the complex population structure, SG Malays have the highest heterozygosity among East and Southeast Asian populations, further confirming their population-genetic diversity (**Figure 2E**).

### Imputation in worldwide populations

Given the genetic diversity represented by the three major ethnic groups in Singapore, we expect our SG10K data (Phase 1) to be a valuable resource for imputation in Asian populations. To assess the effectiveness of our dataset for imputation, we obtained array genotyping data for 53 worldwide populations from the HGDP^26^ and three Singapore populations from the Singapore Genome Diversity Project (SGVP)^21^ and performed imputation experiments using 1KG3, SG10K, and two combined reference panels (**Materials and Methods**). We masked 10% of the Illumina 650K array genotyped SNPs on chromosome 2 for direct assessment of imputation error rate. When imputing these masked SNPs, we found that the SG10K panel outperformed the 1KG3 panel in all East Asian populations except for Japanese (**Figures 3A and 3D**), which is not surprising given that the Japanese ancestry component was barely found in Singapore populations (**Figure 2**). Compared to 1KG3, imputation with SG10K reduced the error rates by 50% when imputing SG Malay and SG Chinese, and by 10% when imputing SG Indian (**Supplementary Table 7**). Beyond Asia, we found that the SG10K panel also improved imputation in Melanesian and Papuan from Oceania, which is likely due to their shared haplotypes with SG Malay. Nevertheless, as expected, SG10K imputation performed worse than 1KG3 imputation in other continental groups, especially in African populations. Surprisingly however, SG10K also performed worse in most Central and South Asian populations. We note that Central and South Asian populations in HGDP are mainly from Pakistan, which historically received substantial gene flow from Central Asia and western Eurasia.^44,45^ These results reflect the limitation of the SG10K panel in capturing the vast population-genetic diversity in Central Asia. For Singapore populations, we further evaluated imputation quality based on the Rsq statistic.^46^ We observed that the SG10K panel substantially improved rare variant imputation, exemplified by a more than two-fold increase in the number of high-quality imputed rare variants (Rsq>0.8, MAF<0.01; **Figure 4**). Compared to SG Chinese and SG Malay, we found that the improvement in SG Indian was smaller. In addition, we noticed that SG10K imputation had higher median Rsq than 1KG3 imputation across all MAF bins, but had a smaller number of high-quality imputed common variants (Rsq>0.8, MAF>0.2) because by sequencing diverse worldwide populations, the 1KG3 panel catalogs a larger number of common variants.

**Figure 3.**
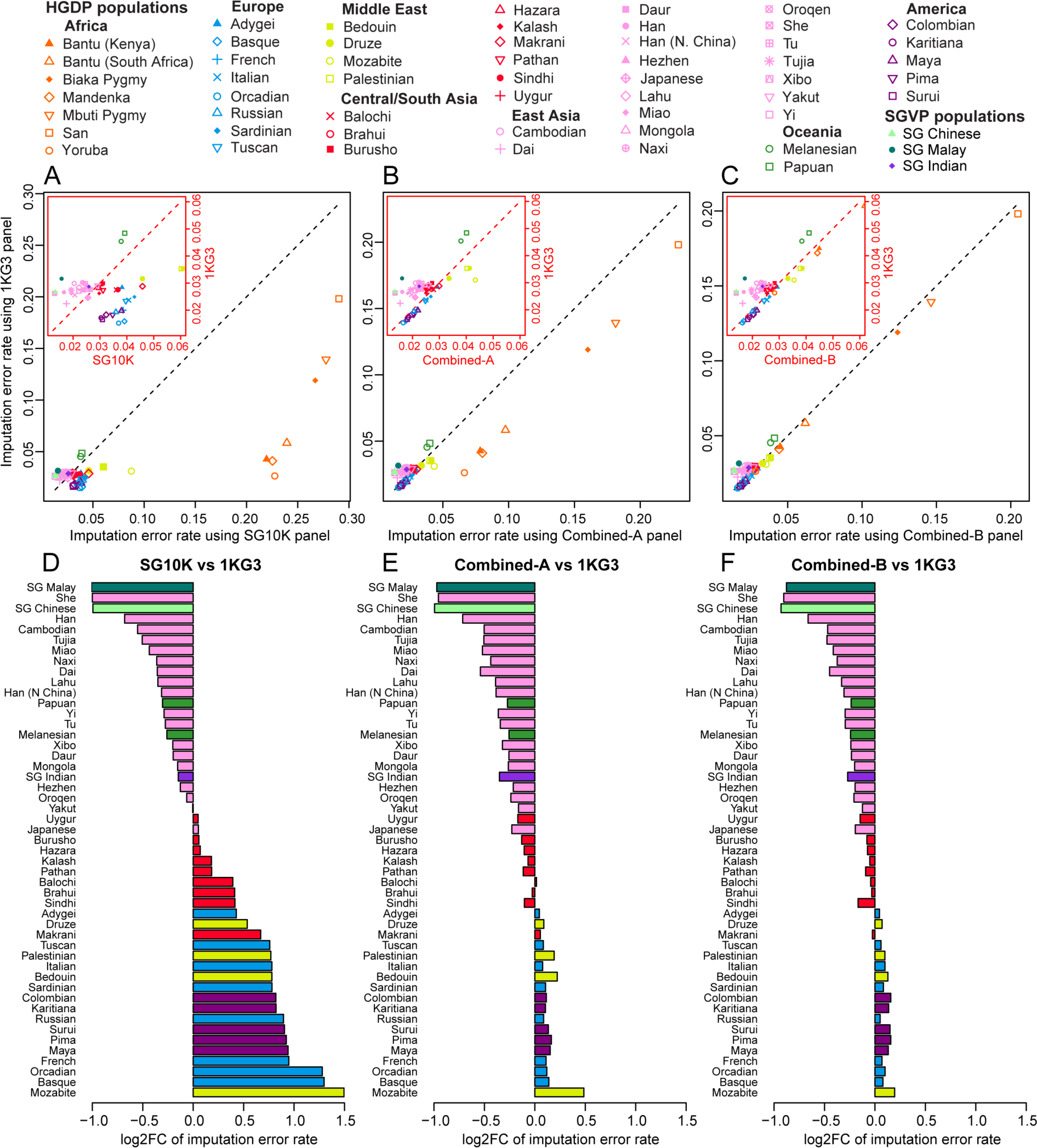
Imputation accuracy in 56 worldwide populations using different reference panels. Four panels were evaluated: 1KG3; SG10K; Combined-A, a merged panel of 1KG3 and SG10K using the reciprocal imputation approach; Combined-B, a merged panel of 1KG3 and SG10K using the naïve reference imputation approach. Imputation error rate was calculated by comparing to the masked genotypes of 4633 SNPs on chromosome 2. (A-C) Imputation error rate using SG10K, Combined-A and Combined-B panels (x-axis) versus error rate using the 1KG3 panel (y-axis). The red inserted boxes are zoom-in plots of the left-bottom corners. (D-F) Fold change (log_2_ scale, log2FC) of imputation error rates for non-African populations using SG10K, Combined-A, and Combined-B panels compared to using the 1KG3 panel.

**Figure 4.**
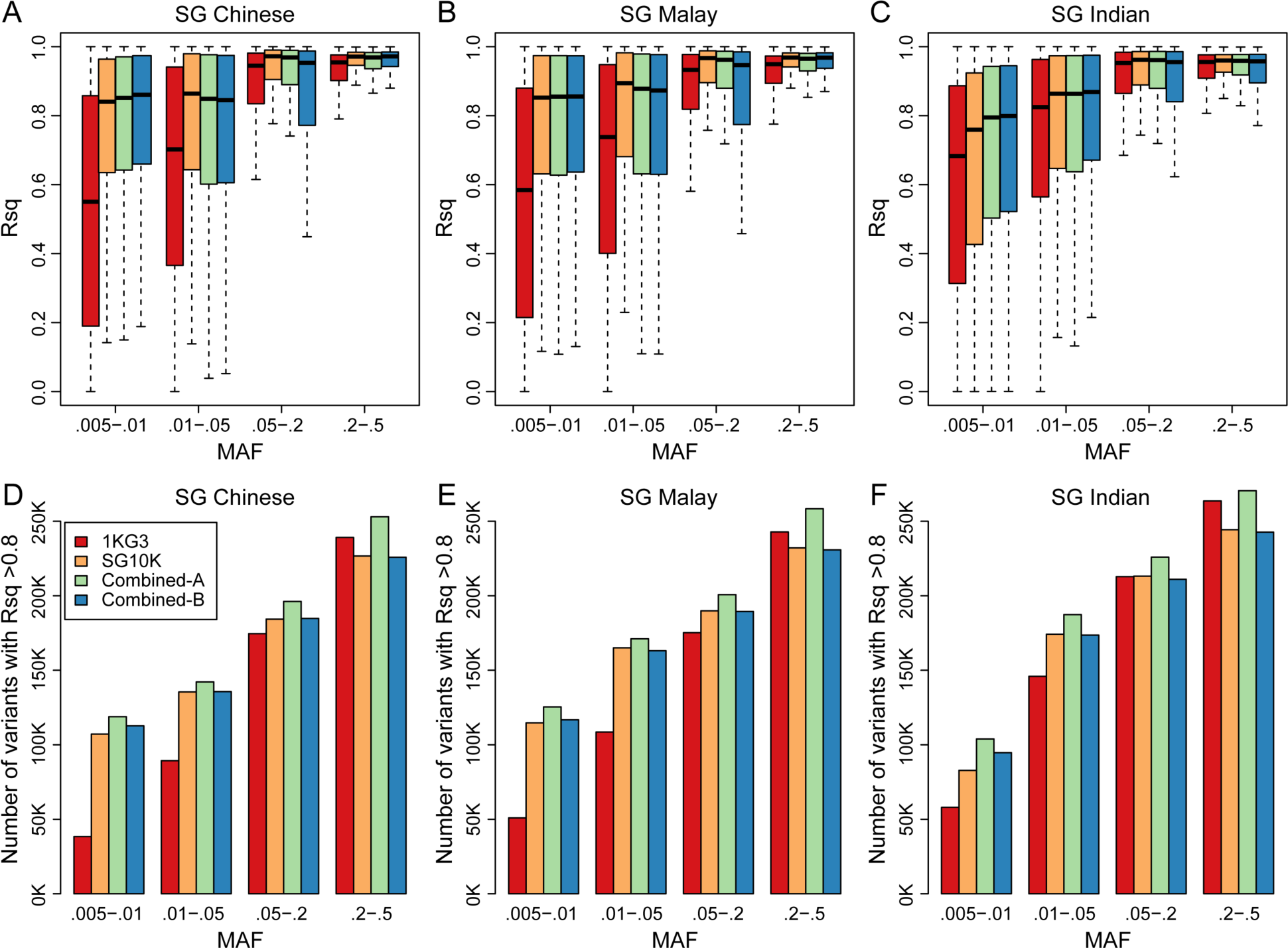
Imputation Rsq in three Singapore populations using different reference panels. The same four reference panels were compared as in **Figure 3**. We evaluated imputation accuracy on chromosome 2 using the Rsq metric calculated for each population from the SGVP dataset. (A-C) Box plots of Rsq values for imputed variants in different MAF bins for each population. Outliers more than 1.5 times of the interquartile range (marked by whiskers) are not shown. (D-F) Number of high-quality (Rsq>0.8) imputed variants in different MAF bins for each population. Different reference panels are coded by colors shown in the legend in (D).

To maximize imputation quality, we sought to create a combined reference panel using SG10K and 1KG3 datasets. This task, however, was not straightforward because ~2/3 of the variants are unique to either SG10K or 1KG3. As a consequence, taking the intersection of these two datasets would lead to a substantial loss of variants. We instead first explored the reciprocal imputation strategy to merge two reference panels to their union set of variants. This strategy used two panels to impute each other reciprocally before merging and was previously proposed to combine the 1KG3 and UK10K datasets for imputation in Europeans.^47^ By adding the SG10K haplotypes, the combined panel (denoted as Combined-A) improved imputation accuracy over the 1KG3 panel in all Asian and Oceanian populations except for a minor decrease in Makrani and Balochi, highlighting the potential broad regional impacts of SG10K (**Figures 3B and 3E**). We found that the imputation error rate for SG Indians is now reduced by 22% compared to when using the 1KG3 panel. For Singapore populations, we observed that the Combined-A panel can impute many more high-quality variants (Rsq>0.8) across all MAF bins than when using either the 1KG3 or SG10K panels alone (**Figure 4**). Nevertheless, we noticed that the Combined-A panel performed much worse than the 1KG3 panel in African populations and Mozabite, even though the 1KG3 haplotypes had been integrated into the panel. This result is because imputing diverse 1KG3 populations with the SG10K as a reference panel might introduce numerous errors in populations distant to the Singapore populations (**Figure 3A**). Furthermore, we found that even for sites that were originally genotyped in 1KG3, LD-based imputation with the SG10K reference panel could mistakenly change the original genotypes of samples from distant populations, such as those from Africa (**Supplementary Figure 18**). We therefore explored an alternative approach to merge the SG10K and 1KG3 panels by imputing sites absent from one dataset as reference homozygotes. This naïve imputation strategy did not change the sites originally genotyped. As expected, we found that the new panel (denoted Combined-B) achieved imputation accuracy comparable to the 1KG3 panel in populations distant to Singapore populations (**Figures 3C and 3F**). However, due to the low accuracy inherent to the naïve imputation approach, the Combined-B panel performed worse than the Combined-A panel for imputing Singapore populations as well as a number of Asian populations (**Figures 3 and 4, Supplementary Table 7**). In summary, our extensive evaluation of the efficacy of different imputation strategies demonstrated the value of SG10K for imputing diverse Asian and Oceanian populations, and highlighted the challenges and potential issues associated with merging two imputation reference panels.

## Discussion

In this article, we have presented a comprehensive WGS-based characterization of genetic variation segregating in 4,810 Singaporeans. In contrast to WGS projects in many countries, which often consist of primarily homogeneous populations, Singapore has diverse populations due to its recent migratory history. Chinese and Indian, the major immigrant populations in Singapore, represent the two largest populations in the world. The population diversity and large sample size together contributed to our discovery of 52 million novel variants. Although 99.5% of the novel variants are rare (MAF<0.01), we identified 126 “deleterious” mutations that are common in Singapore populations but absent from existing public databases. We expect to find many more such variants if we relax the criteria from “absent” to “present at low frequency” in existing public databases. Filtering candidate variants by population allele frequencies is an essential step to pinpoint causal mutations in the diagnosis of Mendelian diseases.^48^ Our findings reiterate the importance of population-specific reference data for reducing genetic misdiagnoses,^11^ and support our key objective to set up a reference database for a long-term national precision medicine program in Singapore.

Using this dataset, we gained novel insights into the geographic and genetic structure of Asian populations. Malay represents indigenous people in Southeast Asia and contributes a novel ancestry component that was not captured by the 1000 Genomes Project.^5,22^ We observed a clear north-south clinal pattern of genetic variation in both South Asia and East/Southeast Asia, except for two recent migrant populations--the SG Chinese and SG Indian, which is consistent with previous studies that suggest a strong role of geography in producing human population structure.^49,50^ Moreover, we found noticeable amounts of admixture among the three major populations in Singapore. In addition, we identified two closely related ancestral components (components 4 and 5 in **Figure 2E**) that are prevalent in East and Southeast Asian populations, suggestive of their ancient origins. Based on the geographic distributions of these two components, we speculate that they might reflect two waves of prehistoric migration from south China to Southeast Asia through a mainland route (component 5) and an island route (component 4). This hypothesis is consistent with a complex peopling history of Southeast Asia depicted by a recent ancient DNA study.^51^ The study suggested that an expansion from East Asia into mainland Southeast Asia occurred about 4,000 years ago during the Neolithic transition to farming, and that an island route migration corresponding to the Austronesian expansion into Philippines and Indonesia took place about 2,000 years ago. However, we were unable to test this hypothesis directly using our data due to lack of samples from other countries in the island Southeast Asia. Future investigation will likely yield a clearer picture as more WGS or dense genotyping data from indigenous populations along the route of the Austronesian expansion become available.^4,52^

In addition to insights about population structure, due to the genetic diversity of Singapore, our project has yielded a key advantage over others in terms of improving genotype imputation in diverse Asian and Oceanian populations. Such improvement is evident when using the combined reference panel of our SG10K data and the 1KG3 data. We note that although it has become a common practice to merge multiple reference panels for better imputation, researchers need to be cautious with using the combined panels.^20,47^ In our case, though the combined panel consisted of worldwide populations from 1KG3, it was not suitable for imputing samples from Africa, Middle East, Europe, and America. As we have demonstrated, this unexpected result was due to errors introduced by the reciprocal imputation procedure to merge two reference panels. The same issue applies to many other studies that attempt to combine population-specific reference panels with global reference panels such as the 1KG3 dataset. We therefore believe that an important future research topic is to study how to optimally combine all published WGS data to generate a mega reference panel that can be applied universally to impute worldwide populations. This endeavor is particularly important for medical genetics, in which reduction of imputation errors can be translated into substantial gain of statistical power in association tests.^53^ Coupled with a substantially improved coverage of rare variants and the rapid accumulation of Asian GWAS data, we expect our SG10K data to be a valuable resource to advance genetic studies of heritable traits and complex diseases in Asian populations, and to mitigate the population disparity in current human genetics research.

## Materials and Methods

### Sample collection

We collected whole blood DNA samples of 5,424 individuals from nine cohorts in Singapore, including 3303 healthy individuals (no major diseases) contributed by the Singapore Eye Research Institute (SERI, N=1,536), Tan Tock Seng Hospital (TTSH, N=971), GUSTO/S-PRESTO birth cohort (GUSTO/S-PRESTO, N=571), SingHealth Duke-NUS Institute of Precision Medicine (PRISM, N=100), the Peranakan Genomes Project (Peranakan, N=79), and the Platinum Asian Genomes Project (Platinum, N=46, consisting of 14 trios and 1 quartet). The remaining 2,121 individuals are patients from three disease studies: heart failure (HF, N=1540), Parkinson disease (PD, N=355), and obese patients undergone bariatric surgery (Bariatric, N=266). These studies were approved by the Institutional Review Board of the National University of Singapore (Approvals: N-17-030E and H-17-049), SingHealth Centralized Institutional Review Board (Approvals: 2006/612/A, 2014/160/A, 2010/196/C, 2009/280/D, 2014/692/D, 2002/008g/A, 2013/605/C, and 2015/2308), National Health Group Domain Specific Review Board (Approvals: 2007/00167, TTSH/2014-00040, 2016/00269, 2018/00301 and 2009/00021). All participants provided written informed consent. However, 571 samples from the GUSTO/S-PRESTO birth cohort were excluded from further analyses after joint genotype calling and phasing due to no consent for data release. Demographic information of samples from the other eight cohorts was summarized in **Supplementary Table 1**. We also obtained province-level ancestry information for 118 Chinese samples from the SERI cohort, whose parents were both from the same province in China. These samples include 35 from Fujian, 54 from Guangdong, 14 from Hainan, and n<3 samples from any other provinces.

### Whole genome sequencing

Genomic DNA was either extracted at the Genome Institute of Singapore (GIS) using QIAamp DNA Blood Midi Kit (Qiagen) or extracted by each contributing study prior to delivery to GIS. DNA quantification was performed by Qubit (Promega) fluorometer with Quant HS dsDNA Assay Kit, High Sensitivity (Invitrogen) followed by 1% GelRed (biotium) stained Hyagarose (Hydrogene) agarose gel electrophoresis run at 100-120 volts for 60 minutes to interrogate DNA integrity using Qubit dsDNA HS Standard #2 DNA (Invitrogen) and 1kb DNA ladder (NEB). Library preparation was undertaken as per protocol using either the Illumina TruSeq Nano DNA, TruSeq DNA PCR-Free or NEBNext Ultra II DNA Library Prep Kit for Illumina (NEB). Paired-end 151bp whole-genome sequencing with an insert size of 350bp was performed using Illumina HiSeq 4000 and X5 platforms. The target depth was 15× for all samples except for 571 samples from the GUSTO/S-PRESTO birth cohort, which were sequenced at 30×. Four samples failed to be sequenced. For the remaining samples, we aligned read pairs to human reference genome GRCh37 using BWA-MEM (v 0.7.12; -M).^54^ PCR duplicates were removed with samblaster (v 0.1.22).^55^ We sorted and merged aligned read pairs from different sequencing lanes using SAMtools (version 1.3),^56^ followed by base quality recalibration using BamUtil recab (v 1.0.13; --maxBaseQual 40). Finally, BAM files were converted to CRAM files with 8 bins of base quality by BamUtil (v 1.0.14; --binQualS 0:2,3:3,4:4,5:5,6:6,7:10,13:20,23:30). After excluding unmapped reads and PCR duplicates, the mean sequencing depth is 13.7× across 4,849 samples targeted at 15×.

### Quality controls before genotype calling

We inferred ancestry, sex, and contamination rate for each sample based on sequencing reads. Ancestry estimation was based on the LASER software,^28,29^ with reference panels from the HGDP dataset^26^ and the SGVP dataset.^21^ The HGDP dataset consists of genotypes across 632,958 autosomal SNPs (Illumina 650K array) for 938 individuals from 53 diverse populations worldwide. The SGVP dataset consists of genotypes across 1,285,226 autosomal SNPs (Illumina 1M and Affymetrix 6.0 arrays) for 96 SG Chinese, 89 SG Malays, and 83 SG Indians. Both datasets were filtered by MAF>0.01.

We first projected our samples into a worldwide ancestry space generated by the top four PCs of the HGDP data (**Supplementary Figure 2A-B**). Three outliers were found: two clustering with Europeans and one close to the Africans. We excluded these outliers and further projected the other samples on a Singapore ancestry map based on the SGVP data (**Supplementary Figure 2C-D**). Given the missing data and potential errors in self-reported ethnicity, we re-assigned ethnicity to each sample using the ancestry coordinates on the SGVP map. As shown on **Supplementary Figure 2C**, the mean PC1 and PC2 coordinates of SGVP Chinese, Malays and Indians form a triangle with individuals from each ethnicity group cluster around one vertex. We divided the map into three sectors by the medians of the triangle, and assigned ethnicity labels to our study samples based on the sectors they were projected to (**Supplementary Figure 2D**). After removing 36 highly contaminated samples, we labeled 2,780 Chinese, 903 Malays, and 1,127 Indians. The overall concordance rate between our inferred ethnicity and self-reported ethnicity is 96.7% (**Supplementary Table 2**).

We inferred sex of each sample based on the ratio of sequencing depths between chromosome X and autosomes, denoted as X/A ratio. Males are expected to have X/A ratio equal to 0.5 and females are expected to have X/A ratio equal to 1. For each sample, we computed the sequencing depth per chromosome using SAMtools,^56^ discarding reads with mapping quality <20 or base quality <20. Samples with a X/A ratio smaller than 0.75 were inferred as males, while the rest were inferred as females (**Supplementary Figure 3**). In total, we inferred 2462 females (51.2%) and 2,348 males (48.8%). The mismatch rate in comparison to self-reported sex is 0.76% (**Supplementary Table 3**).

We used VerifyBamID (v 1.1.2; --precise –maxDepth 100 –minMapQ 20 –minQ 20 – maxQ 100) to assess the level of DNA contamination for each sample by comparing sequence reads to the allele frequencies of the inferred ethnicity group (Chinese, Malay, or Indian), which were computed using the SGVP data.^27^ 36 samples with an estimated contamination rate α>0.05 were removed (**Supplementary Figure 4A**).

### Genotype calling, phasing, and annotation

We performed variant detection and joint genotype calling using the GotCloud pipeline.^30^ We used the version that was used to produce the freeze 3 variant call set for the TOPMed Program (see URLs), adjusting for sample contamination rates. The initial call set consists of 107,293,300 single nucleotide polymorphisms (SNPs) and 15,518,308 insertions and deletions (INDELs) on autosomes, and 4,568,579 SNPs and 686,037 INDELs on X chromosome. Multi-allelic variants were coded as multiple bi-allelic variants. A support vector machine (SVM) classifier was used to filter low-quality variants. We trained the SVM model for autosomes using variants on chromosome 1. Positive labels were defined as known variants genotyped by the 1000 Genomes Project on Illumina Omni2.5M array and negative labels were defined as variants that failed in >2 of the six hard filters on variant features (ABE>0.7, |STZ|>5, IBC<-0.1, IOR>2, CYZ<-5, QUAL<5). Definition of the variant features can be found on the TOPMed pipeline website. For the X chromosome, we excluded the feature of inbreeding coefficient (IBC) from the SVM model and trained the model using X chromosome variants. The SVM filter removed 5,097,496 SNPs (Ts/Tv=1.04) and 4,307,458 INDELs on autosomes and 213,308 SNPs (Ts/Tv=1.00) and 174,871 INDELs on X chromosome. Following SVM, we further filtered variants with excessive heterozygosity (EXHET), defined as sites with a Hardy-Weinberg Equilibrium (HWE) p value <10^-6^ in the direction of excessive heterozygosity. The EXHET filter removed 620,625 SNPs (Ts/Tv=1.03) and 455,285 INDELs on autosomes and 22,973 SNPs (Ts/Tv=0.86) and 18,802 INDELs on X chromosome. The low Ts/Tv ratios of the excluded SNPs suggest the effectiveness of both the SVM filter and the EXHET filter.

To improve genotyping accuracy, we used the BEAGLE software (version 4.1), which took genotype likelihoods as inputs, to perform LD-based genotype refinement.^57^ To speed up computation, we ran BEAGLE in parallel by splitting each chromosome into chunks of 10,000 variants with an overlap of 1,000 variants between neighboring chunks. Splitting and merging were performed using the splitvcf.jar and mergevcf.jar programs in BEAGLE Utilities. Low quality variants with dosage R^2^≤0.3 by BEAGLE were filtered, including 7,314,083 SNPs (Ts/Tv=1.89) and 887,099 INDELs on autosomes and 36,112 SNPs (Ts/Tv=1.55) and 19,201 INDELs on X chromosome.

The genotypes from BEAGLE were used to identify cryptic relatedness and duplicated samples (see next section). Based on the inferred trios/duos and the identified MZ/duplicate pairs, we calculated the total Mendelian and duplicate discordances (DISC) for each variant using PLINK (version 1.9).^58^ We further filtered variants with DISC>2, including 582,277 SNPs (Ts/Tv=0.96) and 509,361 INDELs on autosomes and 30,936 SNPs (Ts/Tv=1.07) and 37,566 INDELs on X chromosome.

We performed population-based haplotype phasing of all 5,381 samples using the default settings of EAGLE (version 2.3.5).^33^ For X chromosome, we first set the heterozygous calls in non-pseudo-autosomal region (non-PAR) for males to missing and then phased all samples together. After phasing, we removed 571 samples from the GUSTO/S-PRESTO birth cohort due to no consent for data release. We also removed 8,290,266 variants that are monomorphic in the remaining samples, including 1,015,402 variants on X chromosome. Proportionally, many more X chromosome variants were removed due to the extra step to set non-PAR heterozygotes to missing before phasing. The final SG10K call set consists of phased haplotypes from 4,810 WGS samples, covering 98,273,706 variants (SNPs and INDELs) from 22 autosomes and the X chromosome. Across 22 autosomes, we have 84,725,366 bi-allelic SNPs (Ts/Tv=2.07), 1,112,853 tri-allelic SNPs, 14,669 quad-allelic SNPs, 3,059,717 insertions and 5,708,237 deletions. On X chromosome, there are 3,275,181 bi-allelic SNPs (Ts/Tv=1.97), 31,853 tri-allelic SNPs, 364 quad-allelic SNPs, 124,180 insertions and 221,286 deletions.

We annotated variants in our final call set using the Ensembl Variant Effect Predictor (VEP) and the corresponding VEP-compiled annotation database (version 91_GRCh37).^59^ Because VEP only annotates deleterious effects predicted by Polyphen^36^ and SIFT^37^ for SNPs, we separately annotated INDELs using the SIFT web server.^38^

### Genetic relatedness and inbreeding coefficient

Based on genotypes from BEAGLE, we used PC-Relate^31^ to estimate both kinship coefficient φ and the probability that two individuals share zero identical by descent (IBD, π_0_).^31^ We focused on the common SNPs (MAF>0.05) overlapping with the SGVP dataset. We aggressively pruned the SNPs to be a least 100 kb apart from each other, resulting in 25,568 SNPs left, in order to accommodate the memory limitation for running PC-Relate on such a large number of samples. We also applied the SEEKIN software (using GT mode and SGVP as reference) to estimate kinship coefficients without pruning SNPs, and obtained similar results.^60^ We classified as *k*-degree related pairs if 2^-k-1.5^< φ< 2^-k-0.5^.^61^ Zero degree means monozygotic twins (MZ) or duplicates. First degree related pairs were further split into parent-offspring (PO) if π_0_<0.1 and full-sibling (FS) if π_0_>0.1. We treated pairs above 3^rd^ degree as unrelated. We used the PRIMUS software to identify duos and trios using the estimated φ and π_0_ between pairs of individuals, as well as age and sex for each individual.^32^ To identify a maximum number of unrelated individuals, we first listed all related pairs, and then removed individuals sequentially from the one that appeared most frequently in the list. Once an individual was removed, the corresponding related pairs were also removed from the list. This procedure iterated until no related pairs remained in the list. The inbreeding coefficient for each sample was estimated as (2φ_ii_-1), where φ_ii_ is the self-kinship coefficient for sample i output by PC-Relate.^31,60^

### Evaluation of genotyping accuracy and sensitivity

We have 1,263 samples from the SERI cohort that were previously genotyped by Illumina Quad610 arrays.^34^ We used the array data across 40,048 SNPs on chromosome 2 to evaluate the quality of our call set. By treating the array data as gold standard, we estimated the sensitivity, non-reference sensitivity, precision, overall genotype concordance, heterozygote concordance, and non-reference concordance of our call set.^62^ Sensitivity is defined as the fraction of polymorphic sites in the array data that were also in the final SG10K call set with both alleles matched. Heterozygote concordance rate is defined as the proportion of concordant genotypes across heterozygous sites in the array data. Definitions of the other four statistics are described in detail by Linderman *et al.* (2014)^62^ and are briefly mentioned in the footnote of **Table 1**. In addition, we designed a novel approach to estimate the sensitivity for detecting rare variants. For all 4,810 samples, we counted the number of non-reference variants with MAF<0.001 and with MAF>0.001 in each sample. We fitted the number of non-reference variants per sample as a linear function of the sequencing depth for each population. The linear model enabled us to estimate the non-reference sensitivity in our call set by assuming 25× WGS achieved 0.99 non-reference sensitivity for detecting variants with MAF<0.001.^24^ We reported the Pearson’s correlation *r* between the number of non-reference variants and the sequencing depth per sample, and tested the null hypothesis of *r*=0 using the Student’s *t* test.

### Principal component analysis, ADMIXTURE, F_ST_, and heterozygosity

We merged our SG10K dataset with the 1KG3 dataset^5^ by extracting 26,748,762 bi-allelic autosomal SNPs called in both datasets, excluding SNPs within five base pairs (bps) of INDELs. We then removed LD by thinning the SNPs to at least 2kb apart using PLINK,^58^ resulting in 1,260,657 SNPs. We investigated population structure using PCA,^29^ ADMIXTURE analysis,^43^ and the F_ST_ and H_e_ statistics.^63^ To avoid the undesirable impacts of oversampling certain populations,^42^ we randomly selected 100 unrelated individuals from each Singapore population and combined with the 1KG3 dataset for PCA. The remaining SG10K samples were projected into the PCA map using LASER.^29^ Analyses were based on 270,909 SNPs with MAF>0.05 in the combined dataset. Similarly, in the analyses of East/Southeast Asians (100 SG Chinese, 100 SG Malays, and 1KG3 East Asians), South Asians (100 SG Indians and 1KG3 South Asians), and Singaporeans (4,446 unrelated SG10K individuals), we filtered SNPs with MAF<0.05 in the corresponding datasets. Based on the first two PCs of East/Southeast Asian analysis and South Asian analysis, we used SVM classifiers to classify SG Chinese and SG Indians into northern and southern groups. The SVM classifiers were trained using CHB and CHS for classifying northern and southern Chinese, and using PJL and STU for classifying northern and southern Indians.

We ran the unsupervised ADMIXTURE analyses on the same set of SNPs as PCA, with the number of ancestral components K from 5 to 15 for the dataset combined with 1KG3 and from 2 to 10 for SG10K dataset alone. For each K, we repeated the analysis at least 7 times with different random seeds and picked the one with the highest likelihood to avoid local minimum. We used the five-fold cross-validation approach in ADMIXTURE to select the optimal K.

We calculated genome-wide F_ST_ between pairs of populations using the Weir-Cockerham estimator.^63^ Hierarchical clustering was applied on the F_ST_ matrix using the complete-linkage method implemented in the *hclust* function in R.

Heterozygosity for each Asian population was calculated by 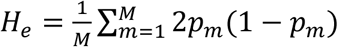 in which *p_m_* is the population-specific allele frequency for the *m*^th^ SNP, and M=237,000 is the number of post-QC SNPs with MAF>0.05 in the combined SG10K and 1KG3 Asian dataset.

### Imputation experiments

We evaluated imputation accuracy both in three Singapore populations from the SGVP dataset, and in 53 worldwide populations from the HGDP dataset. We extracted 46,338 bi-allelic SNPs on chromosome 2, which had consistent alleles in SGVP, HGDP and 1KG3 datasets.^5^ We then masked genotypes of 4,633 SNPs (1 out of every 10 SNPs sorted by position). The masked genotypes were saved for evaluation of imputation accuracy.

We prepared four imputation reference panels for comparison: 1KG3, SG10K, and two combined panels that merge 1KG3 and SG10K datasets. For the 1KG3 panel, we used BCFtools to normalize INDELs, split multi-allelic variants into multiple bi-allelic variants, and removed duplicated variants and extremely rare variants (minor allele count, MAC<5), resulting in a panel of 2,650,510 variants and 5,008 haplotypes (chromosome 2 only). For the SG10K panel, we took the phased data of 4,446 unrelated samples and removed variants with MAC<5, leading to a panel of 2,340,867 variants and 8,892 haplotypes. The number of variants in both 1KG3 and SG10K panels is 1,233,346, indicating a substantial loss of variants if merging by intersection. Instead, we explored two alternative strategies to merge 1KG3 and SG10K to their union set of variants. The first strategy is the reciprocal imputation approach,^47^ in which we used Minimac3 to impute 1KG3 to SG10K and SG10K to 1KG3 respectively,^7^ and then merged the two imputed datasets to form a combined panel (Combined-A). The second strategy is a naïve imputation approach by simply imputing missing data as reference homozygotes using BCFtools, and the combined panel is labelled as Combined-B. After removing 10,111 INDELs that have incompatible allele representations in 1KG3 and SG10K, both combined panels have 3,747,920 variants and 13,900 haplotypes.

Given a reference panel, we pre-phased each populations from HGDP and SGVP using reference-based phasing algorithm in EAGLE2,^33^ followed by imputation using Minimac4.^7^ Imputation error rate was computed for each population as the genotype discordance rate of the 4,633 masked SNPs. In addition, for each of the three SGVP populations, we compared the Rsq statistic for imputed variants in different MAF bins (MAF: 0.005-0.01, 0.01-0.05, 0.05-0.2, and 0.2-0.5).^46^ We did not compare the Rsq statistic for HGDP populations because Rsq cannot be estimated accurately when the sample size of the target population is small.

## Acknowledgements

We would like to thank Hyun Ming Kang, Sayantan Das, Adrian Tan, and Fan Zhang at the University of Michigan and Po-Ru Loh at the Harvard University for helpful discussions. We would also like to acknowledge supports from all the participants and cohort clinical research coordinators. This study was supported by Singapore’s Agency for Science, Technology and Research (Core Funding and Industry Alignment Fund H17/01/a0/007), Biomedical Research Council (Strategic Positioning Fund SPF2014/001), National Medical Research Council (CIRG/1371/2013, CIRG/1417/2015, CIRG/1488/2018, CSA-SI/0012/2017, CG/017/2013, CG/M006/2017_NHCS, NMRC/TCR/013-NNI/2014, STaR/0011/2012, STaR2013/001, STaR/0026/2015, NMRC/TCR/006-NUHS/2013; Centre Grants 2010-13 and 2013-2017),

National Research Foundation (NRF-NRFF2016-03), Core Funding from National University of Singapore, SingHealth and Duke-NUS, Alexandra Health Small Innovative Grant (SIGII/15203), and funding from Tanoto Foundation, Lee Foundation, Boston Scientific Investigator Sponsored Research Program and Bayer. The data management and analysis of the study was also supported by the National Super Computing Center, Singapore.

## Web Resources

TOPMed calling pipeline, https://github.com/statgen/topmed_freeze3_calling

BamUtil, https://genome.sph.umich.edu/wiki/BamUtil

PC-Relate, https://www.rdocumentation.org/packages/GENESIS/versions/2.2.2/topics/pcrelate

PRIMUS, https://primus.gs.washington.edu/primusweb/

SEEKIN, https://github.com/chaolongwang/SEEKIN/

LASER, http://csg.sph.umich.edu/chaolong/LASER/

BEAGLE, https://faculty.washington.edu/browning/beagle/beagle.html

ADMIXTURE, http://www.genetics.ucla.edu/software/admixture/

BCFtools, https://samtools.github.io/bcftools/bcftools.html

SIFT web server, http://sift.bii.a-star.edu.sg/

Variant Effect Predictor, https://asia.ensembl.org/info/docs/tools/vep/index.html

1000 Genomes Project, http://www.internationalgenome.org/

Singapore Genome Variation Project, http://phg.nus.edu.sg/#sgvp

Human Genome Diversity Project, http://www.hagsc.org/hgdp/

gnomAD: https://macarthurlab.org/2017/02/27/the-genome-aggregation-database-gnomad/

## References

1. Welter, D. et al. The NHGRI GWAS Catalog, a curated resource of SNP-trait associations. Nucleic Acids Res 42, D1001–6 (2014).

2. Rosenberg, N.A. et al. Genetic structure of human populations. Science 298, 2381–5 (2002).

3. Garrigan, D. & Hammer, M.F. Reconstructing human origins in the genomic era. Nat Rev Genet 7, 669–80 (2006).

4. The HUGO Pan-Asian SNP Consortium. Mapping human genetic diversity in Asia. Science 326, 1541–5 (2009).

5. The 1000) Genomes Project Consortium. A global reference for human genetic variation. Nature 526, 68–74 (2015).

6. Nikpay, M. et al. A comprehensive 1,000 Genomes-based genome-wide association meta-analysis of coronary artery disease. Nat Genet 47, 1121–1130 (2015).

7. Das, S. et al. Next-generation genotype imputation service and methods. Nat Genet (2016).

8. Ashley, E.A. Towards precision medicine. Nat Rev Genet 17, 507–22 (2016).

9. MacArthur, D.G. et al. Guidelines for investigating causality of sequence variants in human disease. Nature 508, 469–76 (2014).

10. Rehm, H.L. et al. ClinGen--the clinical genome resource. N Engl J Med 372, 2235–42 (2015).

11. Manrai, A.K. et al. Genetic misdiagnoses and the potential for health disparities. N Engl J Med 375, 655–65 (2016).

12. Altman, R.B. PharmGKB: a logical home for knowledge relating genotype to drug response phenotype. Nat Genet 39, 426 (2007).

13. Nelson, M.R. et al. The support of human genetic evidence for approved drug indications. Nat Genet 47, 856–60 (2015).

14. Chatterjee, N., Shi, J. & Garcia-Closas, M. Developing and evaluating polygenic risk prediction models for stratified disease prevention. Nat Rev Genet 17, 392–406 (2016).

15. Hindorff, L.A. et al. Prioritizing diversity in human genomics research. Nat Rev Genet 19, 175–185 (2018).

16. Gudbjartsson, D.F. et al. Large-scale whole-genome sequencing of the Icelandic population. Nat Genet 47, 435–44 (2015).

17. The UK 10 K Consortium. The UK 10K project identifies rare variants in health and disease. Nature 526, 82–90 (2015).

18. Consortium, G.o.t.N. Whole-genome sequence variation, population structure and demographic history of the Dutch population. Nat Genet 46, 818–25 (2014).

19. Collins, F.S. & Varmus, H. A new initiative on precision medicine. N Engl J Med 372, 793–5 (2015).

20. McCarthy, S. et al. A reference panel of 64,976 haplotypes for genotype imputation. Nat Genet 48, 1279–83 (2016).

21. Teo, Y.Y. et al. Singapore Genome Variation Project: a haplotype map of three Southeast Asian populations. Genome Res 19, 2154–62 (2009).

22. Wong, L.P. et al. Deep whole-genome sequencing of 100 southeast Asian Malays. Am J Hum Genet 92, 52–66 (2013).

23. Wong, L.P. et al. Insights into the genetic structure and diversity of 38 South Asian Indians from deep whole-genome sequencing. PLoS Genet 10, e1004377 (2014).

24. Rashkin, S. et al. Optimal sequencing strategies for identifying disease-associated singletons. PLoS Genet 13, e1006811 (2017).

25. Li, Y., Sidore, C., Kang, H.M., Boehnke, M. & Abecasis, G.R. Low-coverage sequencing: implications for design of complex trait association studies. Genome Res 21, 940–51 (2011).

26. Li, J.Z. et al. Worldwide human relationships inferred from genome-wide patterns of variation. Science 319, 1100–4 (2008).

27. Jun, G. et al. Detecting and Estimating Contamination of Human DNA Samples in Sequencing and Array-Based Genotype Data. Am J Hum Genet 91, 839–48 (2012).

28. Wang, C. et al. Ancestry estimation and control of population stratification for sequence-based association studies. Nat Genet 46, 409–15 (2014).

29. Wang, C., Zhan, X., Liang, L., Abecasis, G.R. & Lin, X. Improved ancestry estimation for both genotyping and sequencing data using projection Procrustes analysis and genotype imputation. Am J Hum Genet 96, 926–37 (2015).

30. Jun, G., Wing, M.K., Abecasis, G.R. & Kang, H.M. An efficient and scalable analysis framework for variant extraction and refinement from population-scale DNA sequence data. Genome Res 25, 918–25 (2015).

31. Conomos, M.P., Reiner, A.P., Weir, B.S. & Thornton, T.A. Model-free Estimation of Recent Genetic Relatedness. Am J Hum Genet 98, 127–48 (2016).

32. Staples, J. et al. PRIMUS: rapid reconstruction of pedigrees from genome-wide estimates of identity by descent. Am J Hum Genet 95, 553–64 (2014).

33. Loh, P.R. et al. Reference-based phasing using the Haplotype Reference Consortium panel. Nat Genet 48, 1443–1448 (2016).

34. Cornes, B.K. et al. Identification of four novel variants that influence central corneal thickness in multi-ethnic Asian populations. Hum Mol Genet 21, 437–45 (2012).

35. Tan, A., Abecasis, G.R. & Kang, H.M. Unified representation of genetic variants. Bioinformatics 31, 2202–4 (2015).

36. Adzhubei, I., Jordan, D.M. & Sunyaev, S.R. Predicting functional effect of human missense mutations using PolyPhen-2. Curr Protoc Hum Genet Chapter 7, Unit7 20 (2013).

37. Kumar, P., Henikoff, S. & Ng, P.C. Predicting the effects of coding non-synonymous variants on protein function using the SIFT algorithm. Nat Protoc 4, 1073–81 (2009).

38. Sim, N.L. et al. SIFT web server: predicting effects of amino acid substitutions on proteins. Nucleic Acids Res 40, W452–7 (2012).

39. Landrum, M.J. et al. ClinVar: public archive of relationships among sequence variation and human phenotype. Nucleic Acids Res 42, D980–5 (2014).

40. Yang, S. et al. Sources of discordance among germ-line variant classifications in ClinVar. Genet Med 19, 1118–1126 (2017).

41. Bittles, A.H. & Black, M.L. Evolution in health and medicine Sackler colloquium: Consanguinity, human evolution, and complex diseases. Proc Natl Acad Sci U S A 107 Suppl 1, 1779–86 (2010).

42. McVean, G. A genealogical interpretation of principal components analysis. PLoS Genet 5, e1000686 (2009).

43. Alexander, D.H., Novembre, J. & Lange, K. Fast model-based estimation of ancestry in unrelated individuals. Genome Res 19, 1655–64 (2009).

44. Qamar, R. et al. Y-chromosomal DNA variation in Pakistan. Am J Hum Genet 70, 1107–24 (2002).

45. Majumder, P.P. The human genetic history of South Asia. Curr Biol 20, R184–7 (2010).

46. Li, Y., Willer, C.J., Ding, J., Scheet, P. & Abecasis, G.R. MaCH: using sequence and genotype data to estimate haplotypes and unobserved genotypes. Genet Epidemiol 34, 816–34 (2010).

47. Huang, J. et al. Improved imputation of low-frequency and rare variants using the UK10K haplotype reference panel. Nat Commun 6, 8111 (2015).

48. Whiffin, N. et al. Using high-resolution variant frequencies to empower clinical genome interpretation. Genet Med 19, 1151–1158 (2017).

49. Cavalli-Sforza, L.L.,, Menozzi, P. & Piazza, A.. The History and Geography of Human Genes, (Princeton University Press, Princeton, NJ, 1994).

50. Wang, C., Zöllner, S. & Rosenberg, N.A. A quantitative comparison of the similarity between genes and geography in worldwide human populations. PLoS Genet 8, e1002886 (2012).

51. McColl, H. et al. The prehistoric peopling of Southeast Asia. Science 361, 88–92 (2018).

52. Hudjashov, G. et al. Complex Patterns of Admixture across the Indonesian Archipelago. Mol Biol Evol 34, 2439–2452 (2017).

53. Huang, L., Wang, C. & Rosenberg, N.A. The relationship between imputation error and statistical power in genetic association studies in diverse populations. Am J Hum Genet 85, 692–8 (2009).

54. Li, H. Aligning sequence reads, clone sequences and assembly contigs with BWA-MEM. arXiv: 1303.3997v2[q-bio.GN] (2013).

55. Faust, G.G. & Hall, I.M. SAMBLASTER: fast duplicate marking and structural variant read extraction. Bioinformatics 30, 2503–5 (2014).

56. Li, H. et al. The Sequence Alignment/Map format and SAMtools. Bioinformatics 25, 2078–9 (2009).

57. Browning, B.L. & Yu, Z. Simultaneous genotype calling and haplotype phasing improves genotype accuracy and reduces false-positive associations for genome-wide association studies. Am J Hum Genet 85, 847–61 (2009).

58. Chang, C.C. et al. Second-generation PLINK: rising to the challenge of larger and richer datasets. Gigascience 4, 7 (2015).

59. McLaren, W. et al. The Ensembl Variant Effect Predictor. Genome Biol 17, 122 (2016).

60. Dou, J. et al. Estimation of kinship coefficient in structured and admixed populations using sparse sequencing data. PLoS Genet 13, e1007021 (2017).

61. Manichaikul, A. et al. Robust relationship inference in genome-wide association studies. Bioinformatics 26, 2867–73 (2010).

62. Linderman, M.D. et al. Analytical validation of whole exome and whole genome sequencing for clinical applications. BMC Med Genomics 7, 20 (2014).

63. Weir, B.S. & Cockerham, C.C. Estimating F-statistics for the analysis of population structure. Evolution 38, 1358–1370 (1984).

